# In Depth Flow Inspection Using Dynamic Laser Speckle Temporal Statistics

**DOI:** 10.1101/680330

**Authors:** Mark Golberg, Ran Califa, Sagi Polani, Javier Garcia, Zeev Zalevsky

## Abstract

We present novel optical approach based on statistical analysis of temporal laser speckle patterns for tissue in-depth flow characteristics. An ability to distinguish between Brownian motion of particles and laminar flow is well proved. The main steps in the post processing algorithm and the in-vivo and in-vitro experimental results are presented and demonstrated.

## 1. Introduction

Blood perfusion monitoring is important since tissue blood supply is a key factor in understanding the development of injury and disease. As such, optical non-contact evaluation of tissue blood flow has the potential to provide a dynamic measurement with minimal invasiveness and high spatial resolution [1–3].

When laser light illuminates certain surface, the reflected waves interfere constructively and destructively, resulting in a random interference pattern, widely called a speckle image. Stationary object would create a static speckle image, while a moving object, or object that contains moving particles (as red blood cells), will change the interference pattern, due to the varying phase difference across the illuminating beam. A sequence of speckle frames could be captured and analyzed for motion estimation. Stern et al. [4] were the first to demonstrate the potential of speckle measurement as a tool for blood flow characteristics assessment.

In recent years several methods were adopted as a common practice for estimation of erythrocytes flow velocity through the micro-capillary net in the probed volume. One possible approach is the use of Laser Doppler. The system is usually built of a photodetector and a laser diode, coupled to a pair of spatially separated light transmitting and light receiving fibers. Coherent light that is going through the forward optical channel and reaches the probed tissue volume, is being scattered backwards. As a result, it undergoes Doppler frequency modulation due to the moving elements, such as red blood cells. Some of the frequency modulated light is collected by the receiving optical fibers, induce a random interference pattern. Spectral and/or correlation properties of such pattern depends on the dynamics of the moving particles [5–7].

Another approach is based on the statistical analysis of speckle pattern images created by scattered light. This method is known as the laser speckle contrast analysis (LASCA). Common setup of the LASCA technique constitute from an illumination probe and a statistical analysis of spatial fluctuations of the intensity of speckle modulated images captured with given optics [8–11].

In the above-mentioned methods, there might be a difficulty with distinguishing the exact flow from other in-tissue states, such as increased scatters concentration per a given volume. Our new technique involving usage of speckle statistics and computing the auto-correlation decay model, allows to distinguish between flowing particles and random Brownian motion. In the current paper we developed the theoretical model and demonstrated it in-vitro on a laboratory phantom. Several in-vivo experimental results are presented as well.

## 2. Theoretical Explanation

The proposed method is based on calculating the de-correlation decay time (τ), from the secondary reflected speckle pattern. The data flow comprises of 5 main steps. Figure 1. illustrate the first step being video capturing.

**Figure 1.**
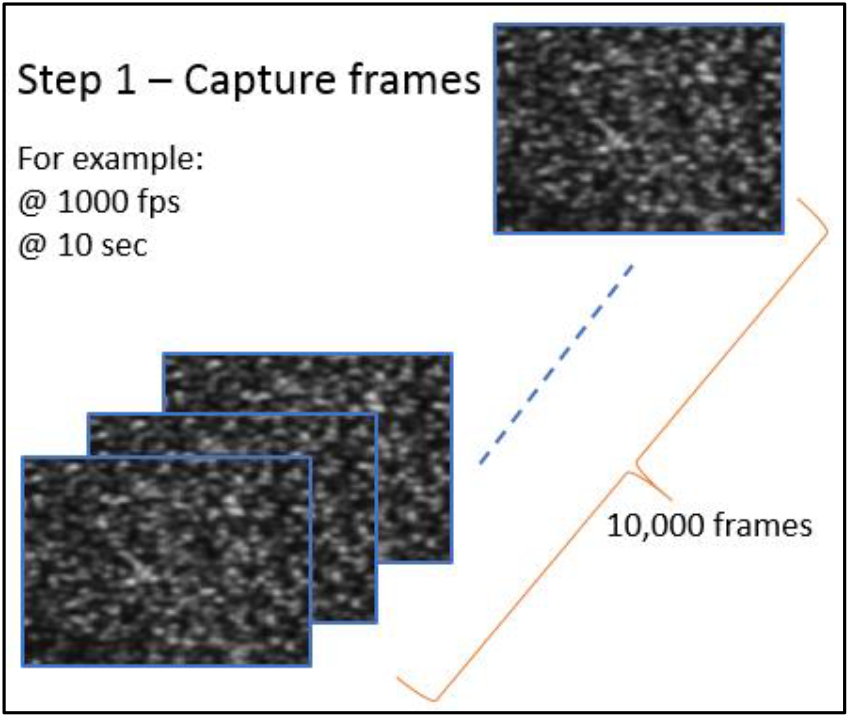
Step 1 – Captured frames.

Step 1: Video of the reflected pattern is being recorded (e.g. 10 seconds of data).

Figure 2 demonstrates video division to “time-wise” batches.

**Figure 2a and Figure 2b.**
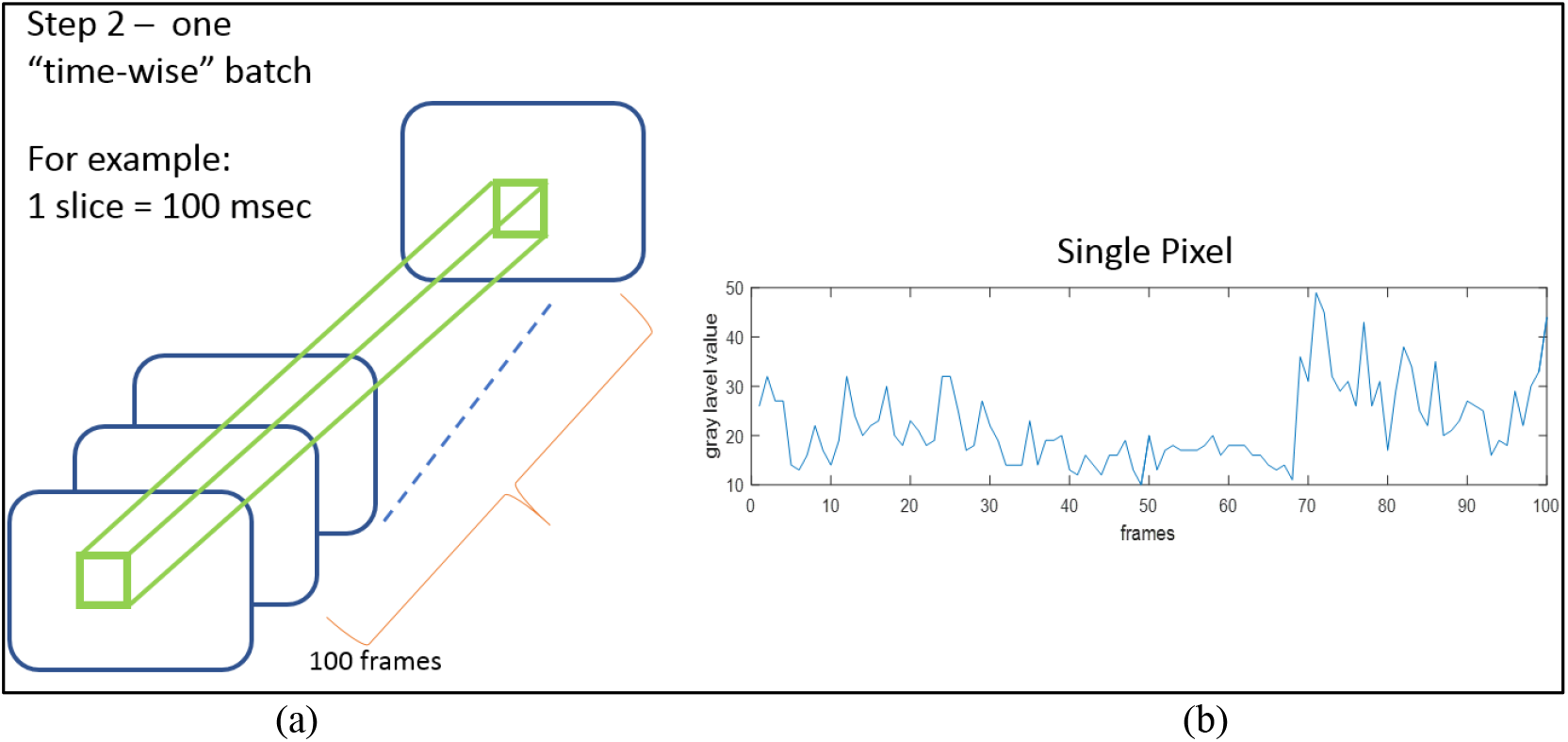
“time-wise” batch. gray level vs. frames of single pixel.

Step 2: Recorded data is divided into time-wise batches (e.g. 100 milli-seconds of data).

Step 3: For each “time-wise” batch, an autocorrelation function of single pixel’s gray levels is being calculated. Auto-correlation function (ACF) is defined by Equation 1 below.

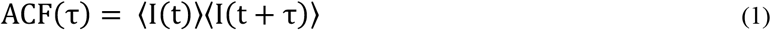

Where I, is the intensity (gray level) per pixel, t is the time and τ is the decay time. Figure 3 below shows the application of Equation 1 on the graph depicted in Figure 2b.

**Figure 3.**
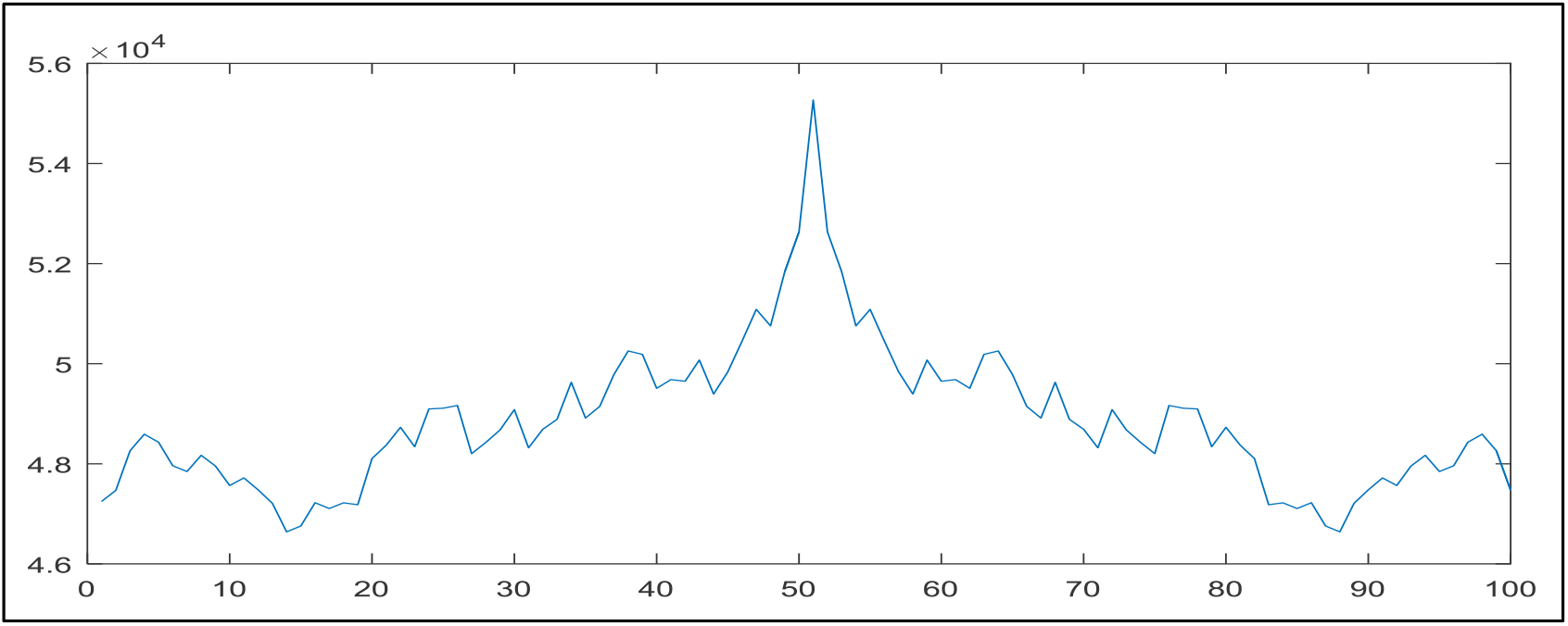
Step 3 – ACF of one pixel.

Step 4: For each decay curve from step 3, τ value is defined as the required time for the autocorrelation distribution to decrease to *e*^−1^ of its maximum value.

Step 5: τ values are being averaged over the full region-of-interest (ROI). Figure 4 below illustrates typical averaged decay curve, after normalization to limits of [0,1].

**Figure 4.**
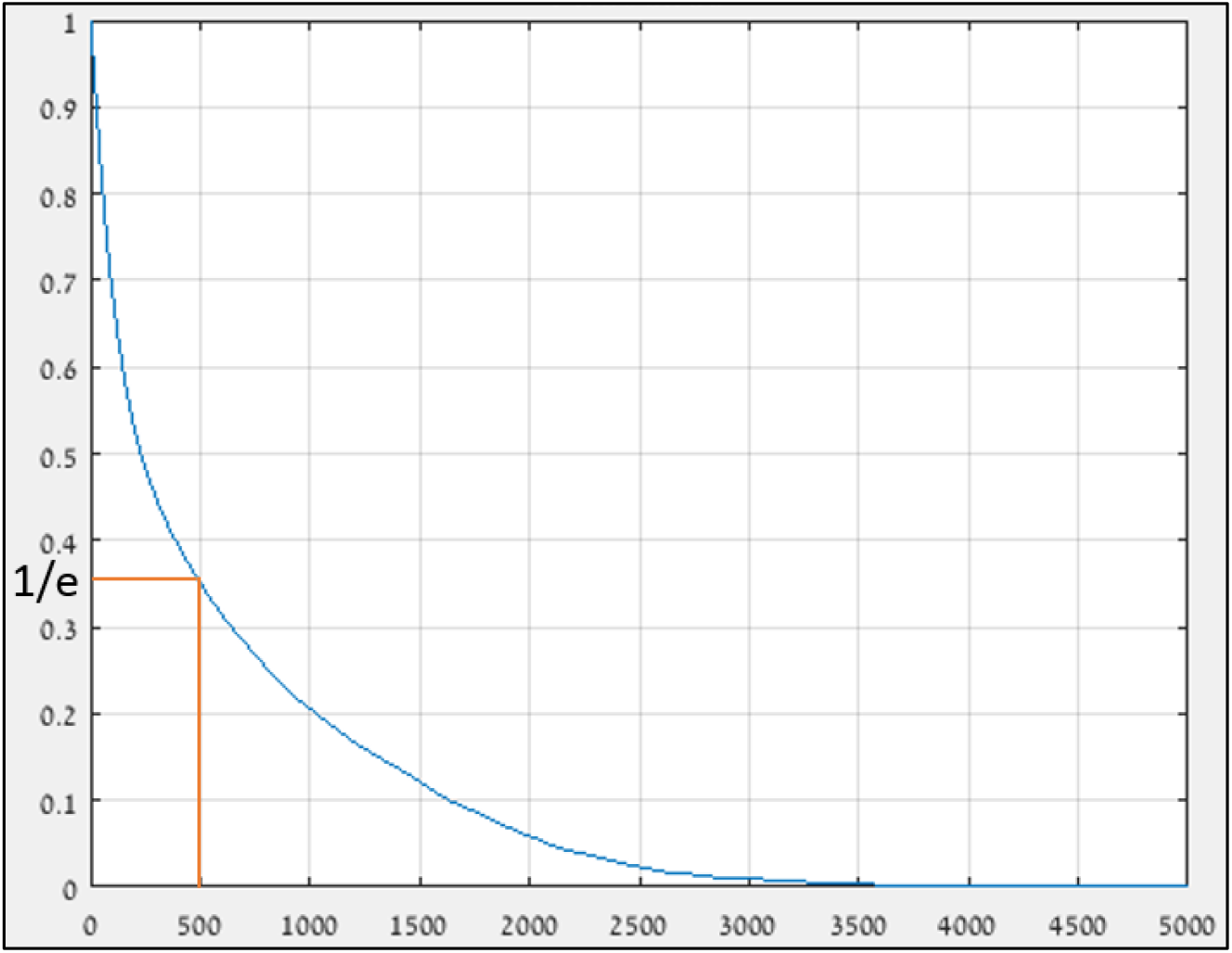
Steps 4 & 5 – typical ACF decay curve

The main innovation in the proposed paper is by modeling the above mention decaying curve. It was shown in previous works [12 – 15], that for moving particles governed mainly by a Brownian motion, a single exponential model (as described in Equation 2 below) could present a high correlation.

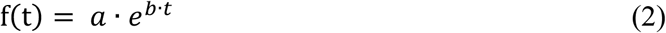

Where, ‘a’ and ‘b’ are specific parameters and t is the time. Alas, the above single exponent model does not work too well for a more complicated motion regime, one that includes flow on top of the Brownian motion. For the latter, flowing particles dynamics, we propose a dual exponent fitting model, as described in Equation 3 below while in section 4, ‘Experimental Results’, we demonstrate its high correlation to model the auto correlation decaying curve.

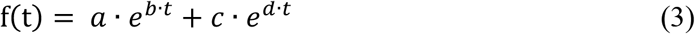

Similarly, to Equation 2. ‘a’, ‘b’, ‘c’ and ‘d’ are specific parameters and t is the time.

## 3. Material and Methods

In this paper we describe several experiments that were done to investigate and demonstrate the theory elaborated in the Section 2. above. The experiments are presented in detail in sections 3.1 – 3.4.

### 3.1. In-vitro setup - flow in a tube – steady state

The following in-vitro experiments were performed with the setup depicted in Figure 5 below.

**Figure 5a and Figure 5b.**
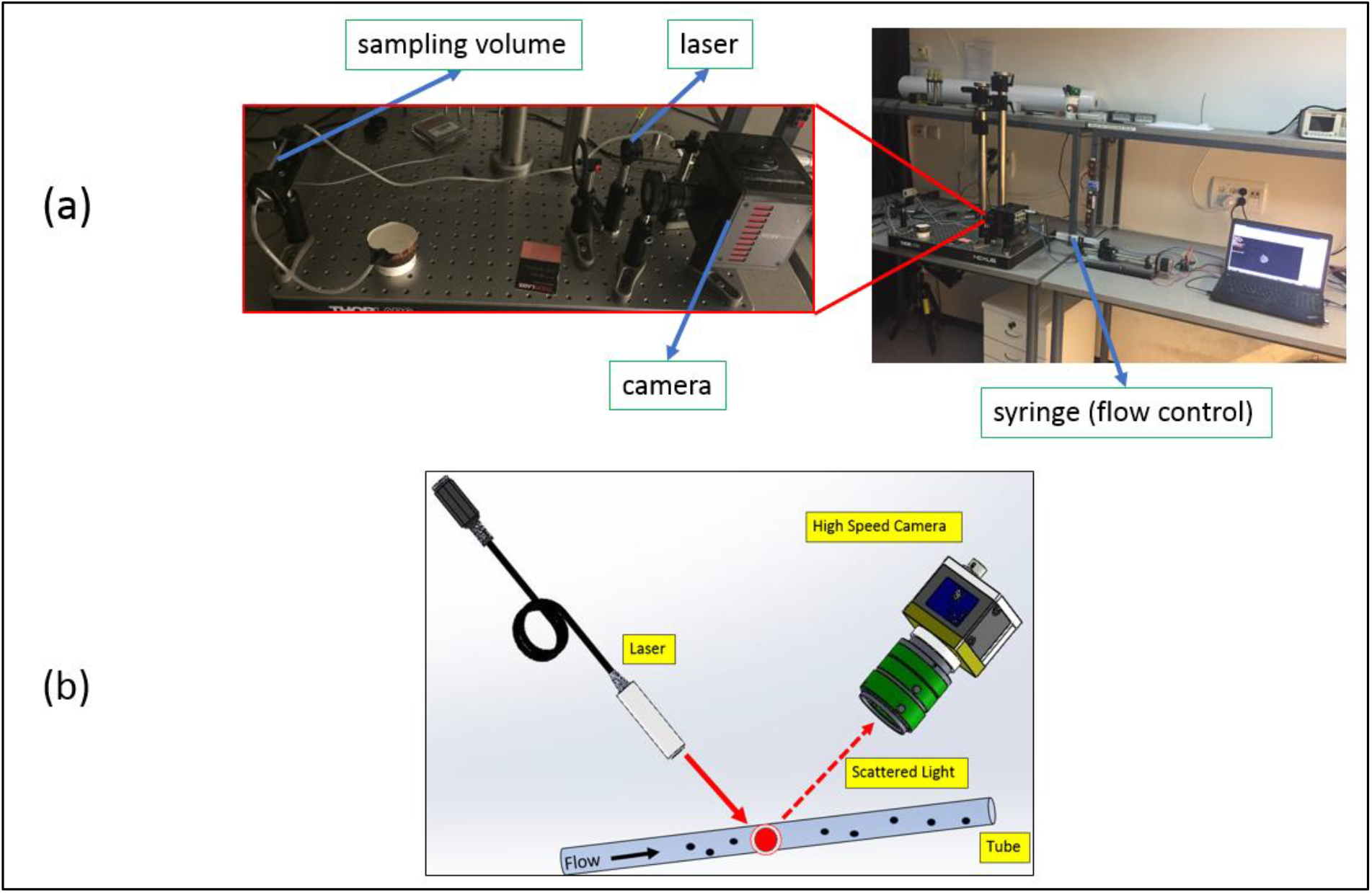
laboratory test setup.; 3D setup illustration.

The optical test setup included a diode-laser (780 nm, 50 mW), a high-speed camera (Fastcam Mini AX200, by Photron) and an objective lens (F_#_2, f35 mm). Tube diameter was 4 mm, with high transparency and made of plastic. A compound of Intra Lipid (0.125%) and Agarose (0.4%) was used as a fluid medium. In section 4 below, one could see the difference in modeling the auto correlation decaying curve, with the two models: one exponent (see Equation 2 above) and two exponents (see Equation 3 above).

### 3.2. In-vitro setup - flow in a tube – transient state

In section 4 below, we present the results of a similar experiment as was described previously section 3.1. The difference is, that flow regime was operated in the following pattern: No Flow --- > With Flow --- > No Flow. The purpose of such configuration was to check the quality of the fit of the dual exponent model in transient states of the flow. For example, when pump is shut down, and flow speed starts to drop.

### 3.3. In-vitro setup - tube diameter identification

The following in-vitro experiment was performed with the setup illustrated in Figure 6 below.

**Figure 6.**
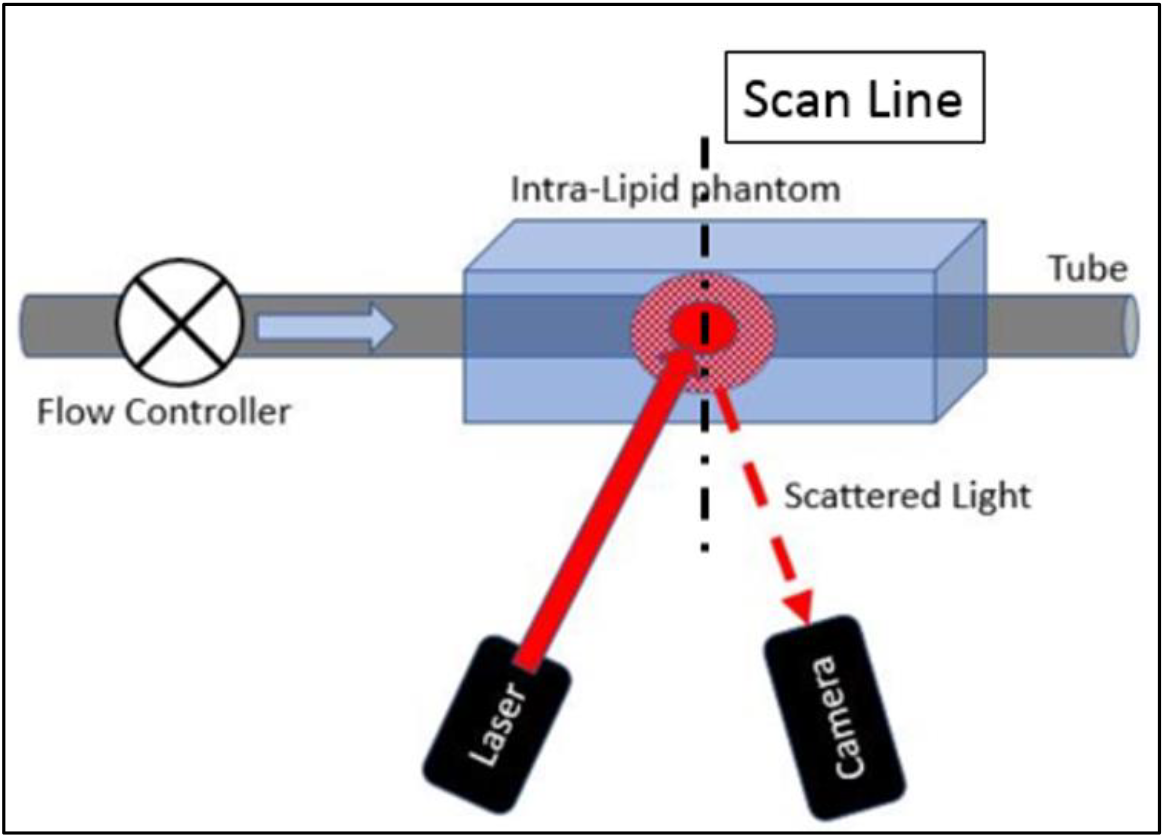
Tube diameter identification – setup illustration.

It’s the same setup as was used in section 3.1 above, only this time we placed the tube inside an Intra Lipid phantom, mimicking an artery within a human tissue (Intra Lipid 0.125%, Agarose 0.4%). Another difference is that instead of illuminating with a single point at a fixed location, this time, we scanned the phantom latterly, by moving the laser illumination spot by 0.5 mm, for each recording session, as could be seen by the black dotted line in Figure 6 above. The capability of the proposed method to identify the boundaries of vessels with flowing particles, “buried” in depth of a scattering medium, is presented in the following section 4.

### 3.4. In-vivo setup - occlusion test

With the same setup as described in Section 3.1 above, an ‘occlusion test’ was performed. With the difference that instead of a tube, the laser illuminated a human tissue. The tip of the index finger was chosen, as an anatomical site with relatively large amount of close to surface blood vessels. Typical sphygmomanometer was used to inflate the cuff to ultra-systolic pressure in order to occlude blood flow. A clear distinguish between the two states: with cuff occlusion and without cuff occlusion, could be seen in section 4 below.

## 4. Experimental Results

As explained in section 3 above, several different tests were performed in order to demonstrate the potential of the method proposed in this paper. Figures 7 – 11 below show the results of those experiments.

**Figure 7a and Figure 7b.**
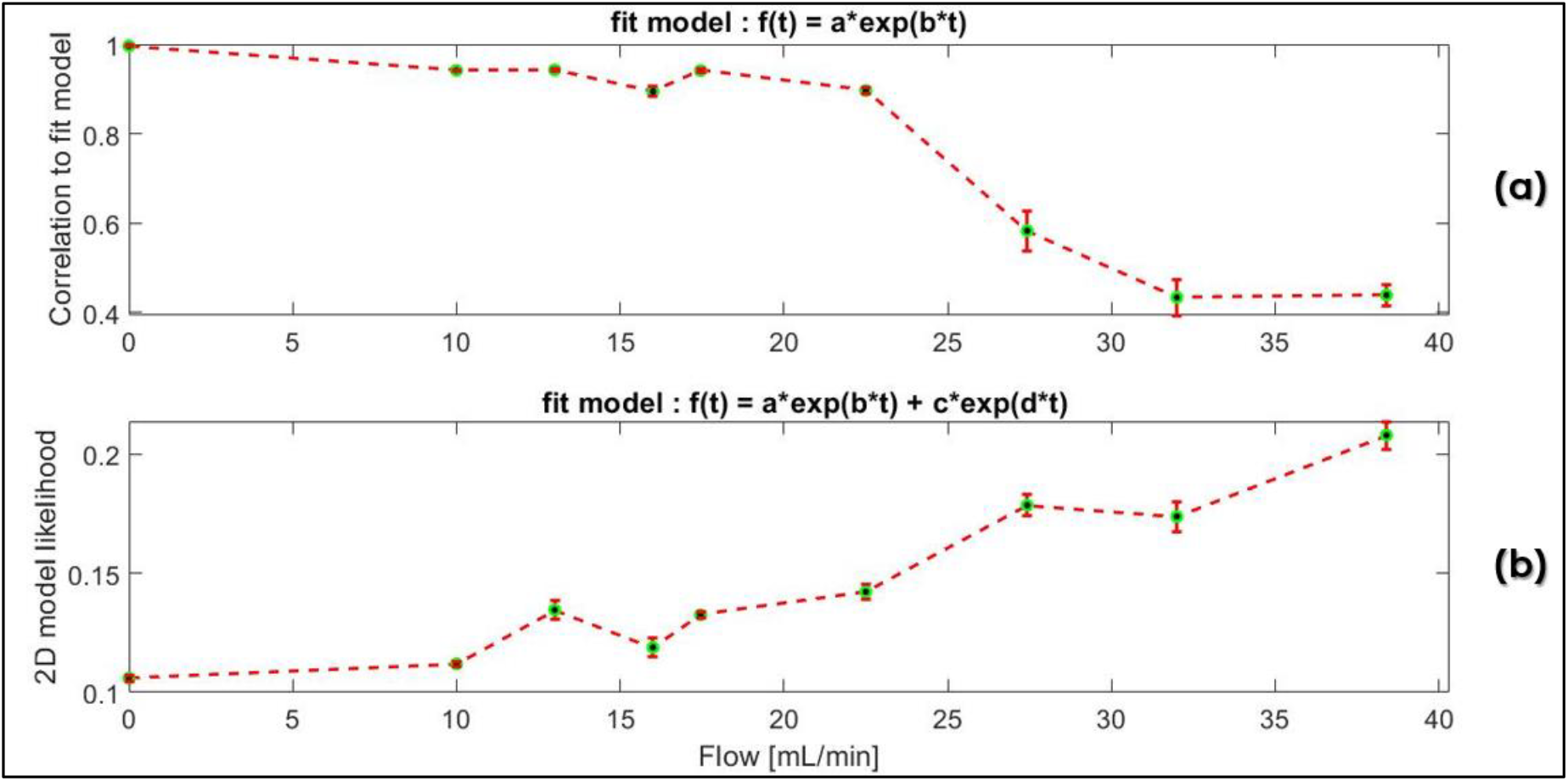
Single exponent model fit.; Dual exponent model fit.

### 4.1 Flow in a tube – steady state

In Figure 7a above, one can see the correlation to a single exponent model, as function of flow speed. As flow increases, the correlation decreases. Such behavior matches our theory, since at high flow rates, particles dynamics is controlled not only by the Brownian motion but also by the force exerted by the pump. Figure 7b demonstrates an opposite trend. This time y-axis is a dimensionless parameter, basically stating how prominent both arguments in Equation 3 above are. It is important to note, that the dual exponent model, can always be reduced to the single exponent model, simply by evoking a very low ratio between the ‘a’ and ‘c’ coefficients. Higher ‘2D model likelihood’ values mean stronger influence of both exponential arguments in Equation 3.

### 4.2 Flow in a tube – transient state

Figure 8 above, shows results of a similar experiment as described in section 4.1, only this time the flow regime was controlled in such a way, that a transient state would occur. Interestingly for us, was to observe the transition from ‘Flow State’ to ‘No Flow State’, as depicted in Figure 8c by a circle. Once pump force is stopped, the flow is decreased to zero, at a rate dictated by the tube properties and by the fluid viscosity. Thus, analysis of model parameters at that specific time, can shine a light on the intrinsic interface between the fluid and its encapsulating vessel.

**Figure 8a, Figure 8b and Figure 8c.**
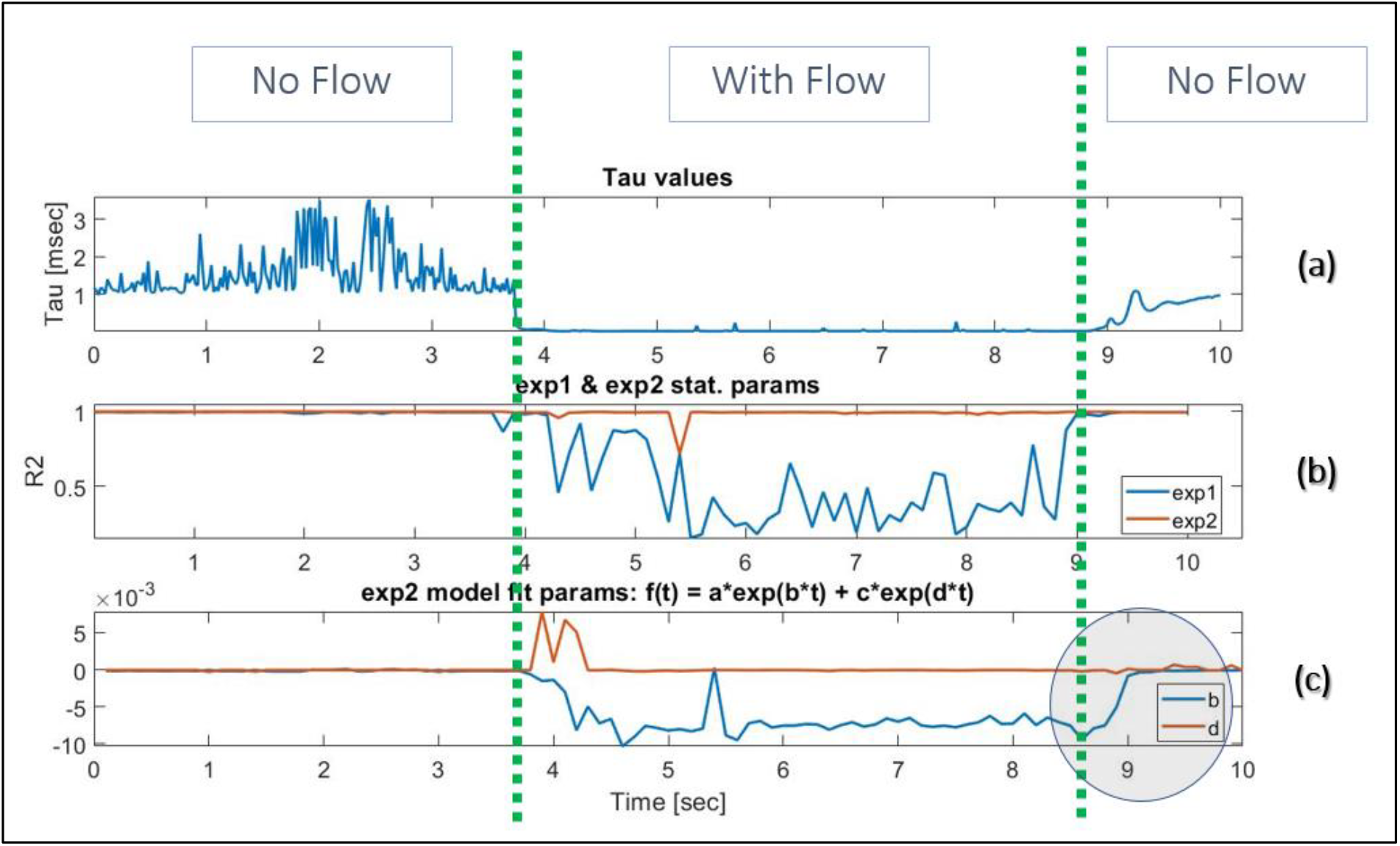
Tau values.; Single & dual exponent models fit.; Dual exponent ‘b’ and ‘d’ parameters.

Figure 8a show the Tau (τ) values. As elaborated in section 2 above, it could be seen that Tau is reciprocal to flow speed. Figure 8b demonstrates once again, the quality of the fit of the dual exponent model vs. the single exponent model. One can clearly see how correlation values drop significantly once flow starts, for the single exponent model (Figure 8b - blue line), while it’s almost constant value of 1, during the entire experiment, for the dual exponent model (Figure 8b - red line).

Figure 9 below shows typical behavior of the ‘b’ parameter from Equation 3 during the transient state. Four different flow rates were tested (Figure 9a-9d), and for each case, the same polynomial fit provided quite high correlation. Such stability in the model parameters, suggests a strong reliability of the model itself.

**Figure 9a – 9d.**
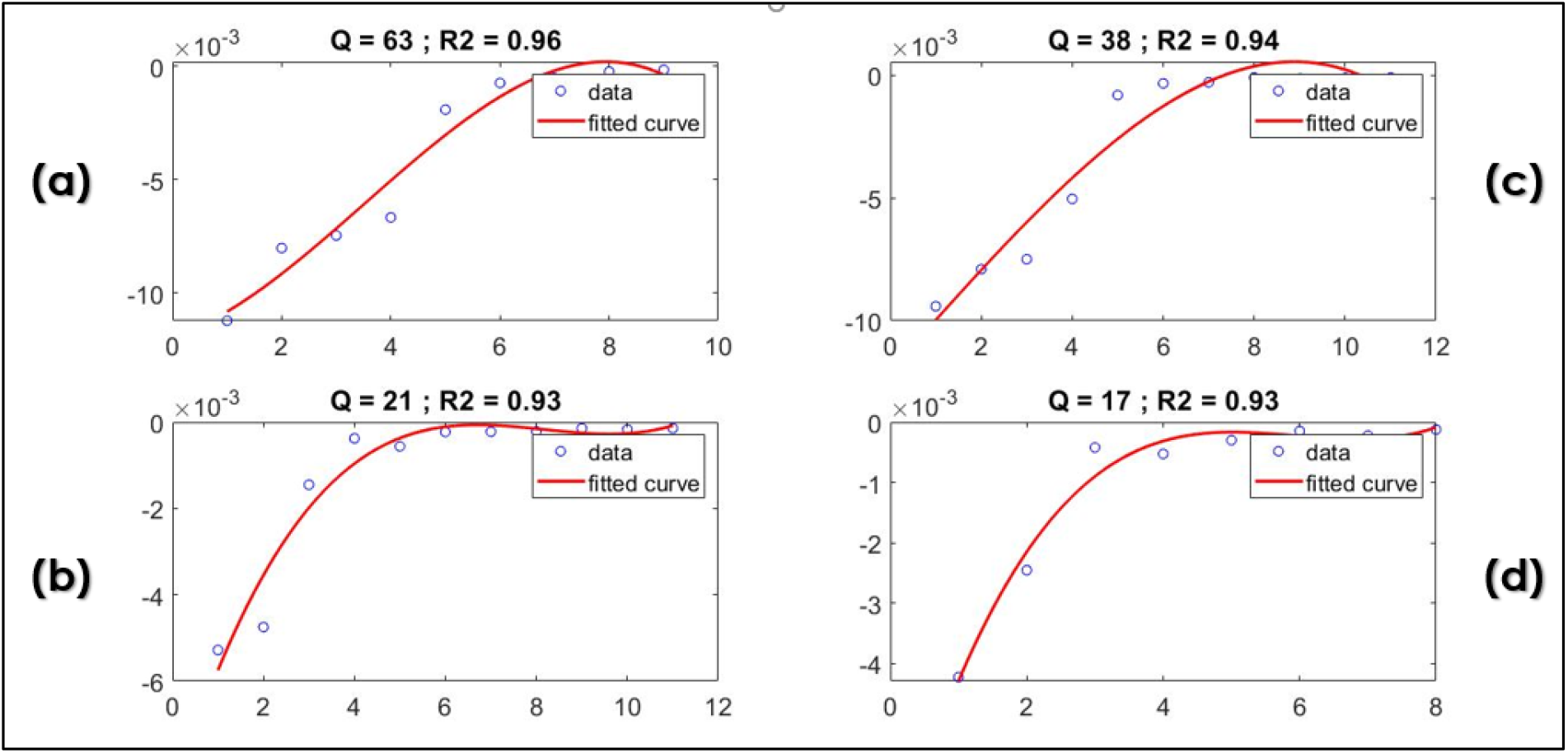
Values of ‘b’ parameter and its corresponding polynomial fit.

### 4.3. Tube diameter identification

In Figure 10 above it could be seen how correlation with the single exponent model (y-axis), changes as a function of the illumination location (x-axis) along the scanning line, as described in section 3.3. Each measurement had a gap of 0.5 mm from its predecessor. Tube outer diameter was 4 mm. Tube was placed in the center of the Intra Lipid phantom. It could be clearly seen that there is an approximate region of 8 steps, at the center of the phantom, where correlation values drop relative of the outer regions. Meaning, in those measurements, the illuminating photons were scattered back to the camera by a flowing particles. As shown in this paper, the single exponent model does not provide a high correlation values for such regime dynamics. By the proposed technique, in depth vessel boundaries could be found.

**Figure 10.**
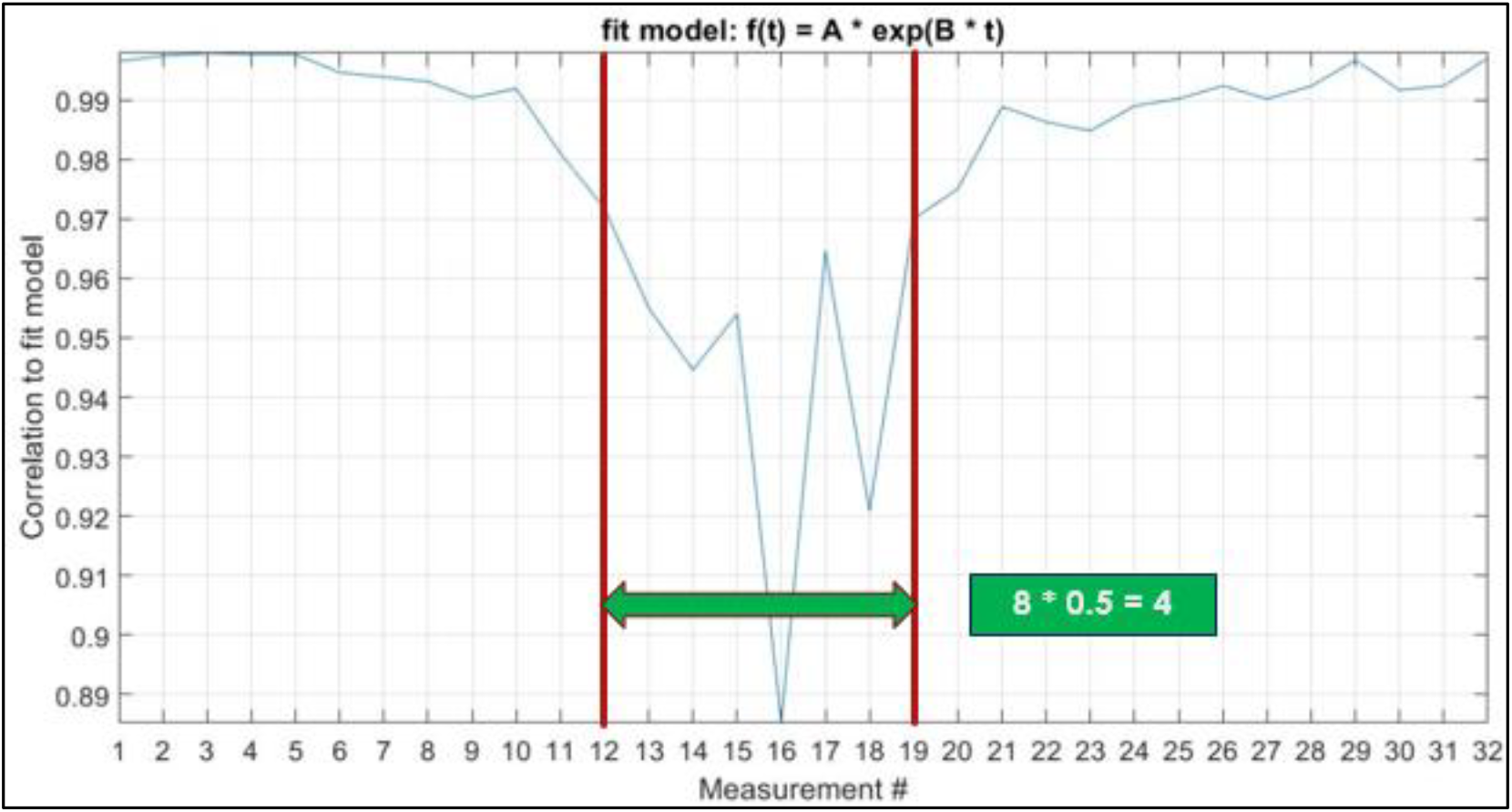
Illustration of the tube diameter identification setup.

### 4.4. Occlusion test

As explained in sections 2 and 3.3 above, when there is a normal blood flow, the correlation to a single exponent model is lower than what occurs in a situation when the cuff is being inflated, and blood flow reduces substantially. See difference between red and blow graphs in Figure 11 above.

**Figure 11.**
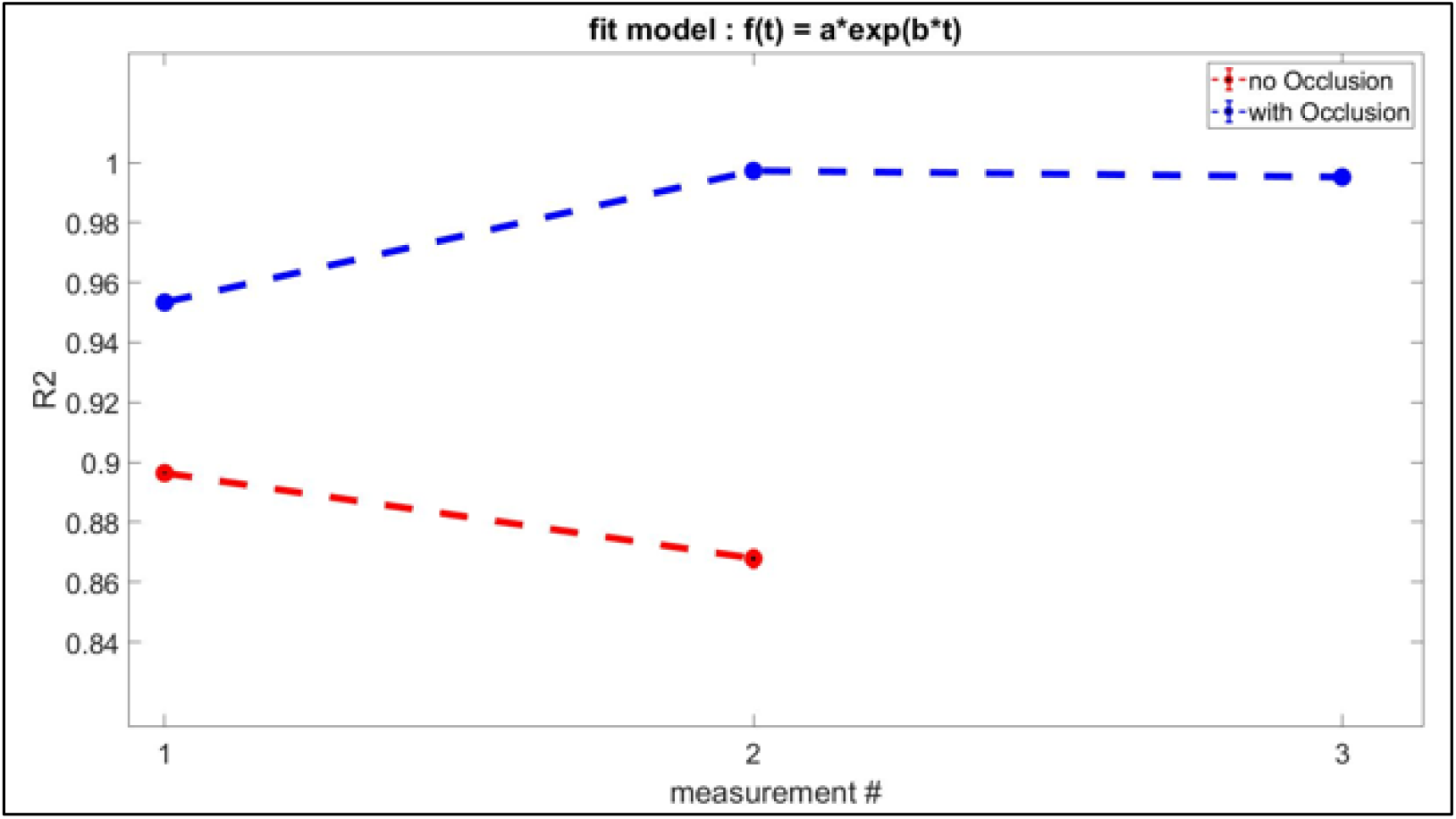
Occlusion test results.

## 5. Conclusions

In this manuscript we presented a novel method for temporal analysis of dynamic speckles generated when a laser beam illuminated a scattering medium such as a tissue. We showed through various experiments, both in vitro and in vivo, the capability of differentiating between Brownian motion and laminar flow motion. We also demonstrated one possible clinical application of identifying the diameter of an in-depth blood vessels. We believe that the above-mentioned analysis could be further investigated to obtain additional physiological parameters (such as, orientation and relative speed of the flow). The main advantage in our method is the relative ease of use, in means of the required hardware complexity and the post processing analytics.

## Acknowledgements

None. No funding to declare.

